# Assessing adverse effects and unspecific effects of transcutaneous spinal direct current stimulation (tsDCS)

**DOI:** 10.1101/2023.12.14.571694

**Authors:** Hongyan Zhao, Ulrike Horn, Melanie Freund, Anna Bujanow, Christopher Gundlach, Gesa Hartwigsen, Falk Eippert

## Abstract

**Background:** Transcutaneous spinal direct current stimulation (tsDCS) is a relatively recent method for non-invasively modulating neuronal activity in the human spinal cord. Despite its growing prominence, comprehensive studies addressing its potential adverse effects (AEs) and unspecific effects (UEs) are lacking.

**Objective:** In this study, we conducted a systematic investigation into the potential AEs and UEs of tsDCS in healthy volunteers.

**Methods:** We used a randomized double-blind within-participant design, employing anodal, cathodal and sham tsDCS of the thoracolumbar spinal cord. Our approach involved a newly-developed structured questionnaire (to assess subjectively-reported AEs) in combination with tsDCS-concurrent recording of skin conductance, cardiac and respiratory activity (to assess UEs in bodily state).

**Results:** The most frequently participant-reported AEs were sensations of burning, tingling, and itching, although they were largely described as mild; skin redness (experimenter-reported) occurred even more frequently. Importantly, when comparing AEs between active and sham tsDCS via frequentist and Bayesian analysis approaches, the results were largely in favour of no difference between conditions (with the exception of skin redness). A similar picture emerged for most UE metrics, suggesting that tsDCS does not induce changes in bodily state, at least as measured by our autonomic nervous system metrics.

**Conclusion:** We believe that the strategy employed here could serve as a starting point for a systematic AE and UE assessment in clinical populations, longitudinal designs and when stimulating different spinal sites. Taken together, our results contribute to assessing the tolerability, safety and specificity of tsDCS, in order to further the investigation of spinal cord function in health and disease.

## 1. Introduction

The spinal cord serves as a hub for the processing and transmission of neural signals between the body and the brain, essential for motor control, somatosensory processing, and autonomic function [1]. Modulating spinal cord function via invasive stimulation has been employed clinically for decades [2, 3], but more recently non-invasive approaches have become feasible as well [4]. Specifically, transcutaneous spinal direct current stimulation (tsDCS) has emerged as a technique for modulating spinal cord excitability [5–9]. Numerous studies have indicated that tsDCS has a modulatory effect on spinal processing related to somatosensory, nociceptive and reflex responses [10–15], suggesting that tsDCS could be a useful tool for investigating spinal cord function in health and disease.

Despite a rapidly growing body of tsDCS studies, the field is lacking systematic studies investigating tsDCS adverse effects (AEs; here defined as subjectively-reported sensations associated with tsDCS) and unspecific effects (UEs; here defined as concurrently-recorded changes in the participants’ physiological bodily state), although such an assessment is important for several reasons. First, it would help to ensure the safety and tolerability of tsDCS by assessing potential risks and discomfort. Second, it would support finding a range of parameter settings that allow for proper blinding, as is of utmost importance especially in clinical settings. And third, being aware of off-target UEs would allow for more informed study design by taking potential confounds into consideration.

Here, we comprehensively assessed possible AEs and UEs induced by tsDCS. First, we performed a systematic keyword search across all published human tsDCS studies to provide an overview of previous work on AE and UE characterization. While such approaches have already been carried out for tDCS [16–18], they are currently lacking for tsDCS. Second, in a preregistered study we performed a detailed questionnaire-based assessment of AEs, including their spatiotemporal properties as well as blinding success. Third, we investigated UEs via tsDCS-concurrent recordings of several physiological parameters to comprehensively assess possible changes in participants’ bodily state. Importantly, both AEs and UEs were assessed in a within-participant design, allowing us to investigate the effects of different stimulation polarities (anodal, cathodal) compared to sham stimulation. Finally, we aimed to not only provide evidence for the possible existence of AEs and UEs, but also for their possible absence (by complementing frequentist analyses with a Bayesian approach [19]), allowing for a rigorous assessment of the safety and tolerability of tsDCS.

## 2. Materials and methods

### 2.1 Assessing adverse effects (AEs) and unspecific effects (UEs) in previous tsDCS work

A systematic literature search was conducted according to PRISMA guidelines [20] across PubMed, Scopus, Web of Science, and Google Scholar to identify human tsDCS studies, using specific search terms for study identification (Supplementary Table 1a), the reporting of AEs (Supplementary Table 1b), and the reporting of UEs (Supplementary Table 1c). Additionally, we explored whether studies reporting positive outcomes in our AE search incorporated questionnaires for AE assessment by examining occurrences of the terms “assessment” and “questionnaire”.

### 2.2 Participants

Twenty healthy volunteers (10 females, mean age: 30.1 years, range: 20-40 years; sample-size specified in a preregistration) participated in this study after providing written informed consent. The study was approved by the ethics committee at the Medical Faculty of Leipzig University.

### 2.3 Experimental design

This study is part of a larger preregistered tsDCS project (ClinicalTrials.gov ID: NCT05711498; OSF-preregistration: https://osf.io/d9tyv; note to preprint readers: the preregistration is currently only available to reviewers). We used a randomized, double-blind, sham-controlled, within-participant design. All participants took part in three sessions, each of which featured a different stimulation condition (anodal, cathodal, sham), with the order being balanced across participants. In order to ensure that participants were aware of the experimental design, the Participant Information Sheet informed them about receiving three different stimulation conditions. Sessions were separated by at least one week (preventing possible carry-over effects from previous sessions), occurred at the same time of day (minimizing effects of diurnal variation) and participants did not take part in other neurostimulation studies during the study (preventing confounding effects).

### 2.4 Transcutaneous spinal direct current stimulation (tsDCS)

tsDCS was carried out using a direct current stimulator (DC-Stimulator Plus, neuroConn, Ilmenau, Germany) with electrodes placed over the thoracic spinal cord (spinous process of the twelfth thoracic vertebra) and the right shoulder (suprascapular region). The target areas were cleaned with alcohol wipes to remove surface grease from skin and thus lower the impedance. We used rectangular rubber-electrodes of 7 x 5 cm size (neuroConn, Ilmenau, Germany) covered with electrode paste (Ten20 Conductive Paste, Weaver and Company, Aurora, USA). Stimulation consisted of a fade-in of 15 seconds, a plateau of 20 minutes (with stimulation at 2.5 mA either anodally or cathodally) and a fade-out of 15 seconds, with tsDCS polarity referring to the electrode placed over the spinal cord. Sham stimulation followed the anodal montage with 15-second fade-in and fade-out, but only 45 seconds of plateau stimulation at 2.5 mA.

### 2.5 Data acquisition

#### 2.5.1 Recording AEs via structured questionnaire

Based on a proposal for a tDCS questionnaire [16], we developed a structured tsDCS questionnaire (Supplementary Figure 1) that allowed us to systematically record i) potential AE symptoms, ii) the relation of reported AEs to tsDCS, iii) participants’ guesses about the authenticity of tsDCS (active or sham; Question 1), iv) participants’ guesses about the direction of tsDCS (inhibitory or excitatory; Question 2), and v) the onset time, duration, and location of reported AEs (Question 3-5). The symptom report part (including Question 3-5) was administered immediately after tsDCS and the questions related to blinding (i.e., Question 1-2) were answered at the end of a session.

#### 2.5.2 Recording UEs via autonomic nervous system measures

During tsDCS, physiological signals were acquired at 2500 Hz using a BrainAmp ExG system (Brain Products GmbH, Gilching, Germany). Skin conductance was recorded by two electrodes placed on the thenar and hypothenar eminence of the right hand, electrocardiographic data were recorded with one electrode placed at the left lower costal arch and referenced to a right sub-clavicular electrode, and respiratory data were recorded via a breathing belt around the lower rib cage.

### 2.6 Data processing

#### 2.6.1 AEs and blinding success

Participants’ ratings of each symptom were scored on a severity scale from 1 to 4 (1: absent, 2: mild, 3: moderate, 4: severe). As these ratings were also used to compute an ‘Aggregate Symptom Score’ (by summing up the ratings across all symptoms), we adjusted them to a scale of 0 to 3, with 0 signifying the absence of AEs in the respective session. Participants’ ratings regarding the relation of symptoms to tsDCS were scored on a scale from 1 to 4 (1: not related, 2: remotely related, 3: probably related, 4: definitely related).

Participants’ answers to questions 1 and 2 were used to assess blinding success, using the following classification: “Active + Inhibitory” was classified as “Anodal“, “Active + Excitatory” was classified as “Cathodal“, and the remaining answers classified as “Sham.” Questions 3 and 4 captured the onset and duration of reported AE symptoms (where participants’ responses were binned into six temporal categories) and question 5 assessed the spatial distribution of AEs (where participants’ responses were binned into four spatial categories).

#### 2.6.2 UEs

All data processing for UEs was carried out using Python 3.9. The summary measures of tsDCS-concurrent physiological signals were extracted for the whole time period of tsDCS and for quarters of that time period.

##### 2.6.2.1 Skin conductance fluctuations (SCF)

Data were down-sampled to 100 Hz and filtered via a bidirectional first-order Butterworth bandpass (passband: 0.0159 Hz to 5 Hz) Spontaneous SCF were quantified via an area under the curve approach, whereby we interpolated over all local minima of the skin conductance time series and determined the area between this baseline signal and the actual time series [21].

##### 2.6.2.2 Electrocardiographic (ECG) activity

R-peaks were automatically detected using a Pan-Tompkins-Algorithm [22] implemented in the python package py-ecg-detectors (https://github.com/berndporr/py-ecg-detectors) and manually corrected. Heart rate (HR) was determined by averaging the heart beats per minute and heart rate variability (HRV) was calculated as the root mean square of successive differences.

##### 2.6.2.3 Respiratory activity

The time points that mark the beginning of a new breathing cycle were automatically detected by determining the signal minima (representing maximum inhalation). Breathing rate (BR) was determined as the number of breaths per minute and breathing rate variability (BRV) was assessed as the standard deviation of the interval between consecutive breaths.

### 2.7 Statistical analysis

As specified in a preregistered analysis plan, we mostly employed one-tailed tests and established statistical significance at a level of p < 0.05. In addition to frequentist tests, we also employed a Bayesian approach by comparing the evidence for the null model against alternative models using Bayes Factors (BF), which allowed us to determine evidence for the presence or absence of an effect [19]. All analyses were carried out in JASP (JASP Team, 2023; version 0.17.3.0; using default uninformed priors), separately for anodal vs sham and cathodal vs sham.

#### 2.7.1 AEs and blinding success

To assess condition differences in AEs, each item of the tsDCS AE symptom report was analyzed separately using a Wilcoxon sign-rank test and the same analysis was carried out on the Aggregate Symptom Score. The participants’ guesses regarding the stimulation condition were analyzed with a McNemar test (not available in Bayesian implementation). The Aggregate Symptom Scores in correctly vs. incorrectly guessing participants were compared with a Mann-Whitney U test.

#### 2.7.2 UEs

The analysis of physiological data was complicated by the fact that in some cases, participants had inadvertently not been instructed not to talk and move during tsDCS administration, leading to abnormal signal fluctuations in these participants’ autonomic measures and the exclusion of several participants’ data (Supplementary Table 2). We compared SCF, HR, HRV, BR, and BRV values between anodal and sham as well as between cathodal and sham. Overall effects (assessing the entire stimulation window) were investigated using paired-samples t-tests and time-dependent effects (quarters of the stimulation window, about 5 minutes each) were investigated using a 2×4 repeated-measures ANOVA, comparing anodal vs sham and cathodal vs sham separately (necessary due to the uneven distribution of missing data mentioned above).

The BF reported for the paired-samples t-tests (BF_10_) indicate the likelihood ratio of the observed data given the alternative hypothesis that the two measures are different in comparison to the null hypothesis that the values are equal. For example, a BF of 3 means that the data are three times more likely to be observed under the alternative than the null and a BF of 1/3 means that the data are three times more likely to be observed under the null than the alternative (conventionally described as providing moderate evidence for the presence or absence of an effect [19].

For the repeated-measures ANOVA, we were interested in the interaction effect of condition and time and report two BF. BF_10_ indicates the likelihood ratio of the observed data given the alternative model that includes the two main factors and the interaction in comparison to the null model that does not contain these elements. BF_incl_ indicates the likelihood ratio of the observed data given models that include the interaction term in comparison to the models that do not include the interaction term.

## 3. Results

### 3.1 Assessing AEs and UEs in previous work

We identified 76 human tsDCS studies, of which 17 did not report any AE search terms (Supplementary Table 3), 14 mentioned at least one search term, but did not observe AEs (Supplementary Table 4), and 45 reported AEs (Table 1). Among the latter, tingling was reported in 33 studies, itching in 22 studies, and burning in 14 studies, with lesser reports of skin-related irritations / sensations, skin redness, and discomfort. An AE assessment based on questionnaires was only carried out in 9 studies and the level of reported details was rather limited (Table 2). A keyword search for UE reporting revealed hits in 10 studies [7, 26–34], but with the exception of one study [23] (which assessed polarity-dependent changes in spontaneous breathing patterns) none obtained tsDCS-concurrent recordings without potential confounds, i.e., the relevant measures were primarily used as an indicator to ensure adequate task performance.

**Table 1.**
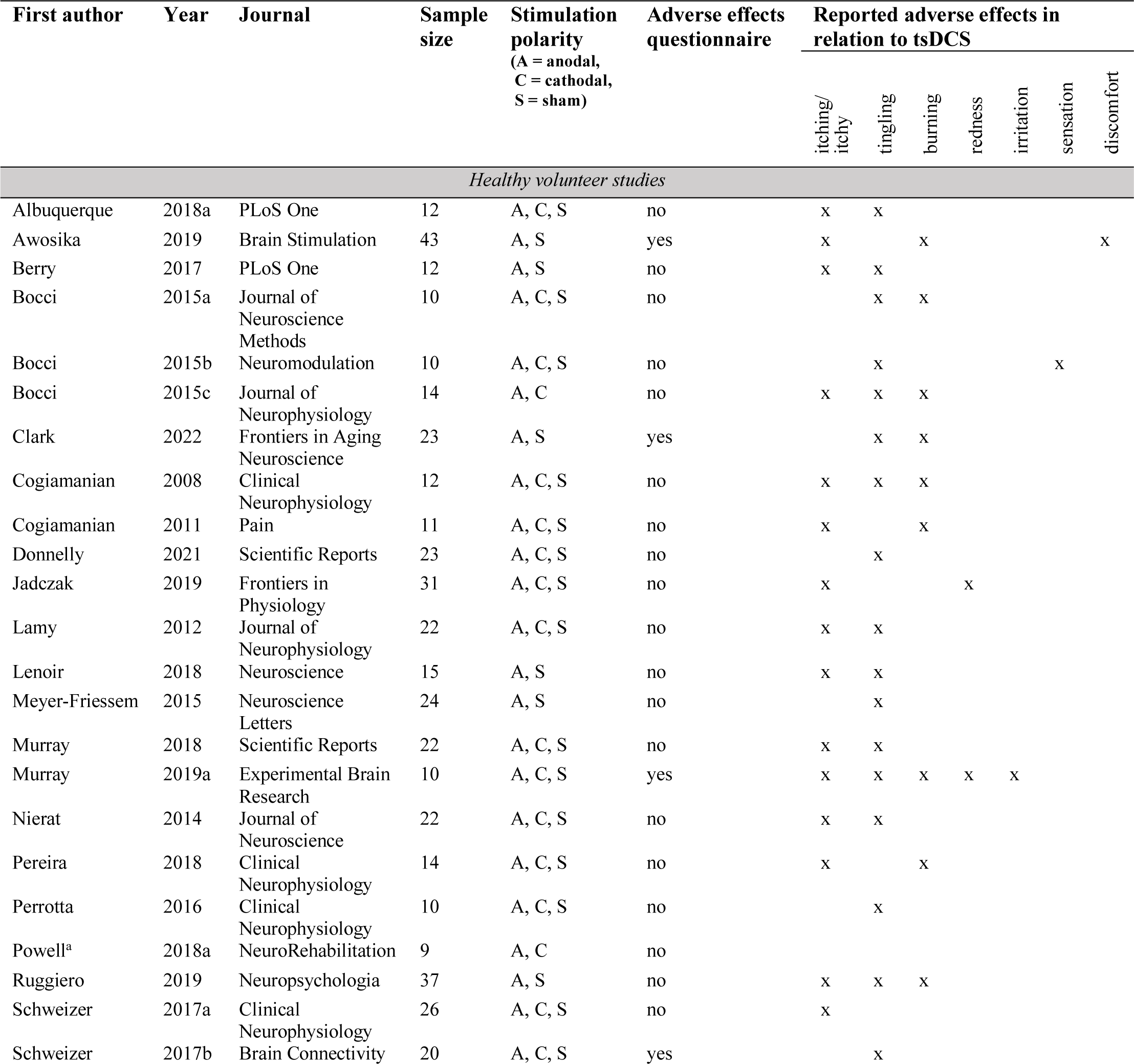
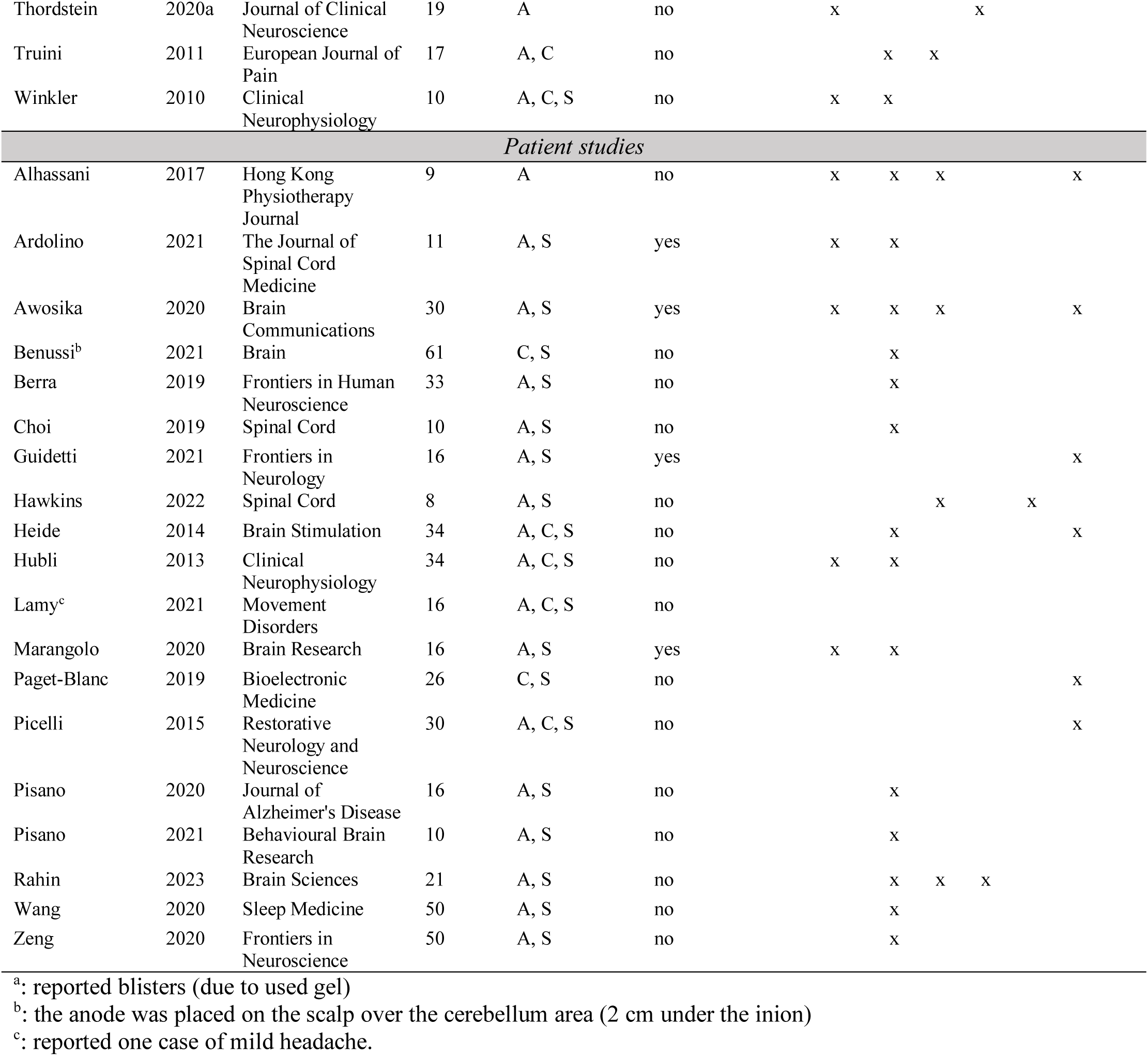
Adverse effects reporting in previous work. This table provides details for all studies in which our systematic keyword search for tsDCS AEs returned hits.

**Table 2.** Questionnaire-based adverse effects assessment in previous work. This table provides details for all studies that used a questionnaire to assess possible AEs of tsDCS

| First author | Year | Journal | Sample size | Stimulation polarity (A = anodal, C = cathodal, S = sham) | Questionnaire for tsDCS adverse effects | Verbatim report of questionnaire results |
| --- | --- | --- | --- | --- | --- | --- |
| *Healthy volunteer studies* |  |  |  |  |  |  |
| Awosika | 2019 | Brain Stimulation | 43 | A, S | Tolerability, Activity and Safety Questionnaire | “Participants’ reports on verbal 0/10 scales indicated the following. In the anodal and sham groups, general discomfort was 1.27 (range 0-5) and 0.90 (range 0-4), perception of pain was 0.18 (range 0-2) and 0.24 (range 0-3), sensation of burning under the electrode was 0.50 (range 0-2), and 0.43 (range 0-5), and itching under the electrode was 0.63 (range 0-2) and 0.76 (range 0-5) respectively. No skin irritation or burns occurred. Thus, tsDCS was overall well tolerated.” |
| Clark | 2022 | Frontiers in Aging Neuroscience | 23 | A, S | “Participants used an 11-point rating scale where 0 represents “none” and 10 represents “strongest/worse possible.” For tsDCS, the following items were rated: tingling, itching, burning, pain, fatigue, nervousness, headache, muscle spasms, mood change, urinary urgency, abdominal/pelvic sensations, and sweating.” | “no adverse effects” ... “For tsDCS there were reports of very mild tingling/burning sensation at the electrode sites (average rating less than 1 out of 10). All other potential side effects of tsDCS were negligible or completely absent.” |
| Murray | 2019a | Experimental Brain Research | 10 | A, C, S | A tsDCS questionnaire was administered “to establish the presence of any adverse effects”, but no further specification was provided. | “Following tsDCS, the major complaint was skin redness or irritation which subsided within a few hours, followed by reports of tingling, burning or itchy sensations mainly during the ramp-up and down phase of stimulation.” |
| Schweizer | 2017b | Brain Connectivity | 20 | A, C, S | „After each session, subjects completed a questionnaire to assess any pain associated with tsDCS as well as their guess as to which polarity of tsDCS they had received.”<br>There was no further specification of the questionnaire. | “No adverse effects from the tsDCS electrodes [...] were reported.” |
| *Patient studies* |  |  |  |  |  |  |
| Ardolino | 2021 | The Journal of Spinal Cord Medicine | 11 | A, S | tsDCS Adverse Effects Questionnaire (Brunoni et al., 2011) | “In general, the experimental procedures were well tolerated by all subjects and only a few subjects reported an occasionally slight tingling or itching sensation beneath the electrodes. No difference was distinguishable between the “real” or “sham” stimulation nor between polarities in relation to sensations caused by stimulation (e.g., itching, tingling, or auditory perception).” |
| Awosika | 2020 | Brain Communications | 30 | A, S | Tolerability, Activity, and Safety Questionnaire | The authors report mean, median and standard deviation of each questionnaire item for anodal and sham stimulation in their Table 2, as well as p-values for the stimulation condition comparison (none of which were significant) |
| Guidetti | 2021 | Frontiers in Neurology | 16 | A, S | tsDCS Adverse Effects Questionnaire (Brunoni et al., 2011) | “No adverse effects were reported” |
| Hodaj | 2023 | Brain Communications | 36 | A, S | Comfort Rating Questionnaire (Palm et al., 2014) | “No adverse effects were reported during or following any of the three interventions.” |
| Marangolo | 2020 | Brain Research | 16 | A, S | Sensation Questionnaire (Fertonani et al., 2010) | “No adverse sensations were reported. Participants did not recognize which condition they were in and they did not detect a difference in sensations between stimulation conditions (Paired sample t-tests: itchiness: $t_{(15)}=-0,19$, $p=0,85$; pain: $t_{(15)}=-0,37$, $p=0,72$; burning: $t_{(15)}=0,24$, $p=0,82$; warmth/heat: $t_{(15)}=-0,27$, $p=0,79$; pinching: $t_{(15)}=-0,37$, $p=0,72$ ; fatigue: $t_{(15)}=0,17$, $p=0,87$).” |

<sup>a</sup>: Please note that the study by Hodaj and colleagues is not listed in Table 1, but in Supplementary Table 4, since these authors did not report AEs.

### 3.2 Assessing AEs

#### 3.2.1 Symptom reports

Turning to our own study, when aggregating data across all conditions in terms of participant-reported symptoms (Figure 1), burning (40.0%), tingling (26.7%), and itching (20.0%) were the predominant AEs (mostly of mild severity), with skin redness (60%) being reported by the experimenter and having the highest occurrence overall and other AEs being virtually non-existent across all 60 sessions.

**Figure 1.**
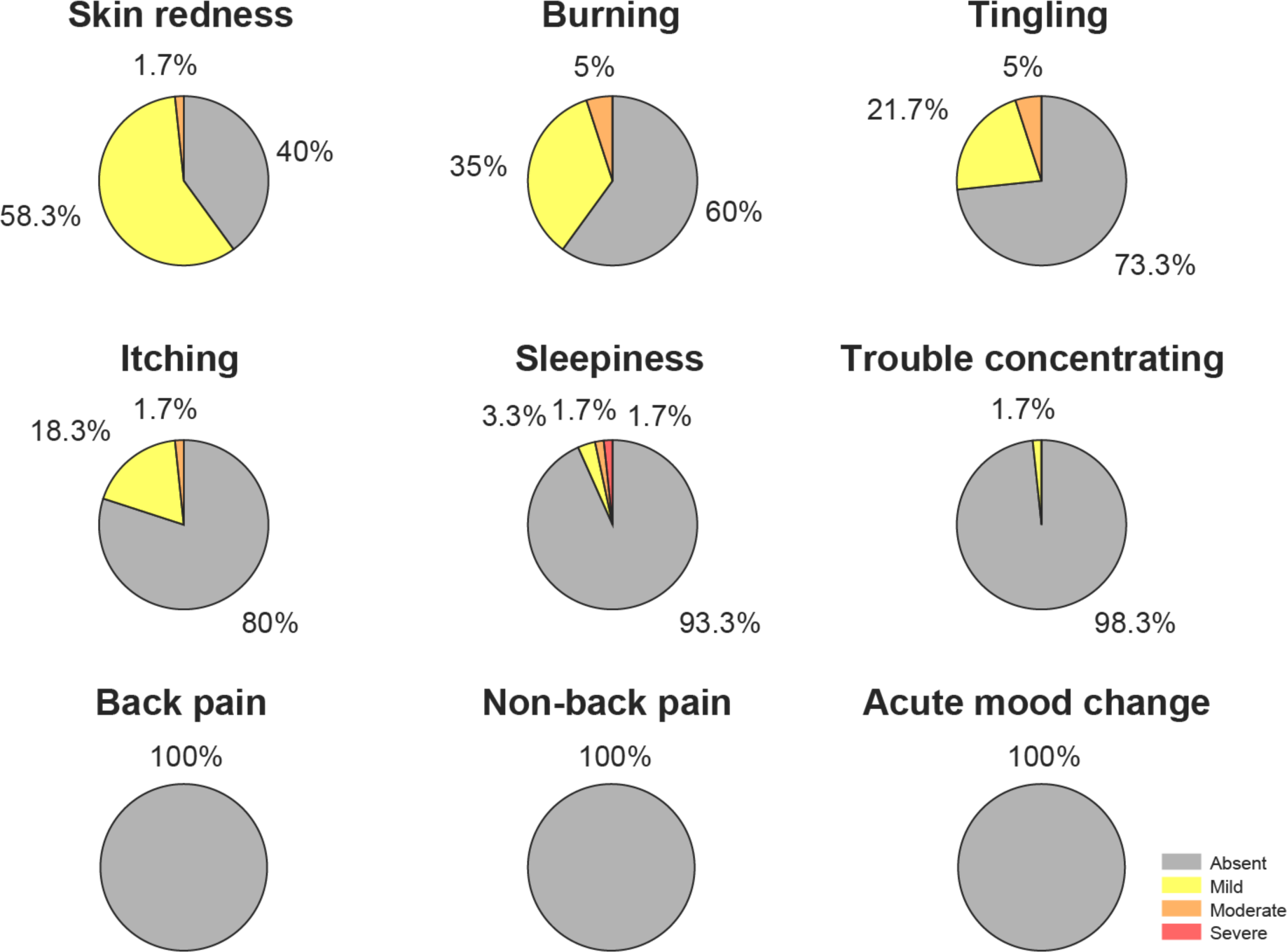
Adverse effects reports. The occurrence and severity of AEs is based on all 60 sessions, with colors representing the severity of the reported adverse effects (see legend).

None of the participant-reported symptoms showed significant differences between conditions and only the experimenter-reported item of skin redness exhibited strong evidence for a difference between the active and sham conditions (Figure 2; Table 3). From a Bayesian perspective, the results clearly favoured the null-hypothesis of no condition differences in participant-reported symptoms over the alternative hypothesis (7/8 BF < 1, 5/8 BF < 1/3 and 0/8 BF > 3). As for the Aggregate Symptom Score, no significant differences were observed for Anodal vs Sham (with the BF being supportive of a null effect), but for Cathodal vs Sham a marginally significant effect was observed, though not paralleled by the BF analysis, indicating inconclusive evidence.

**Figure 2.**
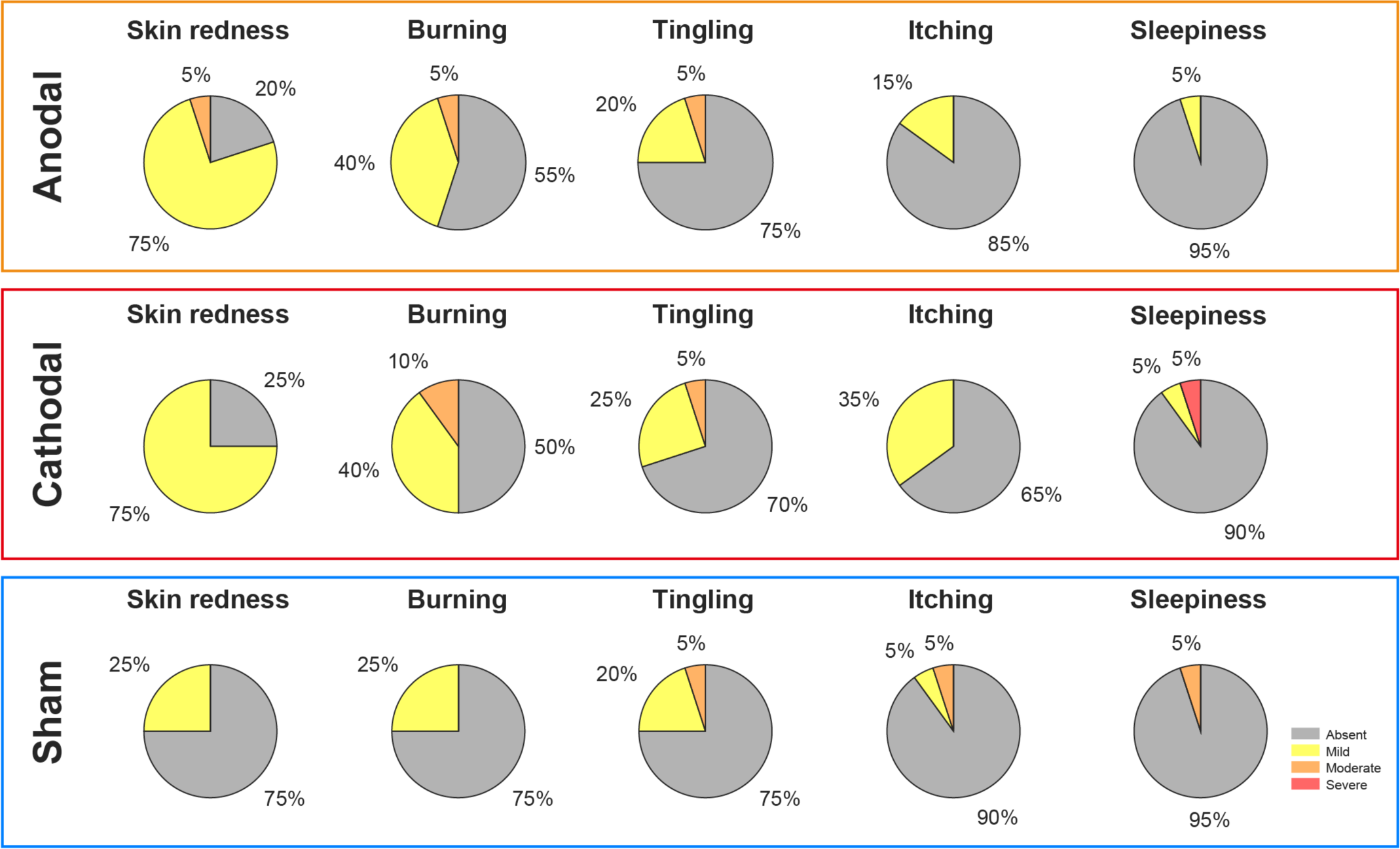
Adverse effects report across conditions. The occurrence and severity of AEs is depicted dependent on condition (Anodal, Cathodal, Sham; each based on 20 participants), with colors representing the severity of the reported adverse effects (see legend). Please note that back pain, non-back pain, acute mood change, and trouble concentrating are not displayed here due to the absence of reports (except for one report of trouble concentrating in the Cathodal group).

**Table 3.** Statistical Comparison of AEs.

| Comparisons | Anodal vs Sham |  | Cathodal vs Sham |  |
| --- | --- | --- | --- | --- |
|  | Wilcoxon signed-rank test |  | Wilcoxon signed-rank test |  |
| Reported AEs | p value | BF<sub>10</sub> | p value | BF<sub>10</sub> |
| Skin redness | 0.002 | 42.07 | 0.002 | 22.45 |
| Tingling | 1 | 0.27 | 1 | 0.29 |
| Itching | 1 | 0.27 | 0.29 | 0.48 |
| Burning | 0.19 | 0.58 | 0.06 | 1.45 |
| Sleepiness | 1 | 0.30 | 0.75 | 0.31 |
| Back-pain | --- | --- | --- | --- |
| Non-back pain | --- | --- | --- | --- |
| Trouble concentrating | --- | --- | --- | --- |
| Acute mood change | --- | --- | --- | --- |

| Comparisons | Anodal vs Sham |  | Cathodal vs Sham |  |
| --- | --- | --- | --- | --- |
|  | p value | BF<sub>10</sub> | p value | BF<sub>10</sub> |
| Symptom Score | 0.43 | 0.30 | 0.042 | 1.53 |

Note that the symptom score does not include skin redness, i.e., only aggregates participant-reported symptoms.

Regarding the reported relation between AEs and tsDCS, skin redness, tingling, itching, and burning were reported as highly associated with tsDCS, while sparsely reported symptoms exhibited a much weaker reported relationship with tsDCS (Supplementary Figure 2).

#### 3.2.2 Assessing participant blinding

When assessing participants’ assumptions regarding the type of stimulation they had received (“Active” or “Sham”), 5% indicated they had received 0/3 active sessions, 20% thought 1/3 were active, 50% indicated that 2/3 were active, and 25% believed 3/3 were active. Upon assessing participants’ reports regarding the specific stimulation type they had received, 5% of participants had entirely incorrect answers, 55% had one correct answer, 35% had two correct answers, and only 5% had entirely correct answers. When testing if participants were able to correctly guess the stimulation condition, we observed no significant effects (anodal vs sham: p = 0.74; cathodal vs sham: p = 0.62). Based on the Aggregate Symptom Score, we also explored if participants’ subjective symptom experiences were related to the accuracy of their guesses, but found no evidence for this: all p > 0.4 and all BF < 0.6.

#### 3.2.3 Assessing temporo-spatial AE properties

The reported AEs exhibited distinct patterns in terms of onset time, duration, and location across the different stimulation conditions (Figure 3). In the sham condition, no AEs were observed in half of the participants and the onset of the reported AEs mostly occurred during the tsDCS initiation phase and all within the first 5 minutes (Figure 3A). In the active conditions, AE onset showed a clear shift towards later onset times compared to the sham condition. With respect to the duration of AEs (Figure 3B), all reported AEs for sham stimulation occurred within the initial 5 minutes, whereas reported AEs for active stimulation conditions had a much longer duration. Most AEs were reported to occur at the back electrode site in both active and sham stimulations. This was followed by reports of occurrence under both electrodes, yet here more prominently in active compared to sham conditions (Figure 3C). Experimenter-reported skin redness was notably absent in the majority of participants (75%) during sham stimulation, contrasting with active stimulation, where it predominantly occurred at the shoulder electrode site (Figure 3D).

**Figure 3.**
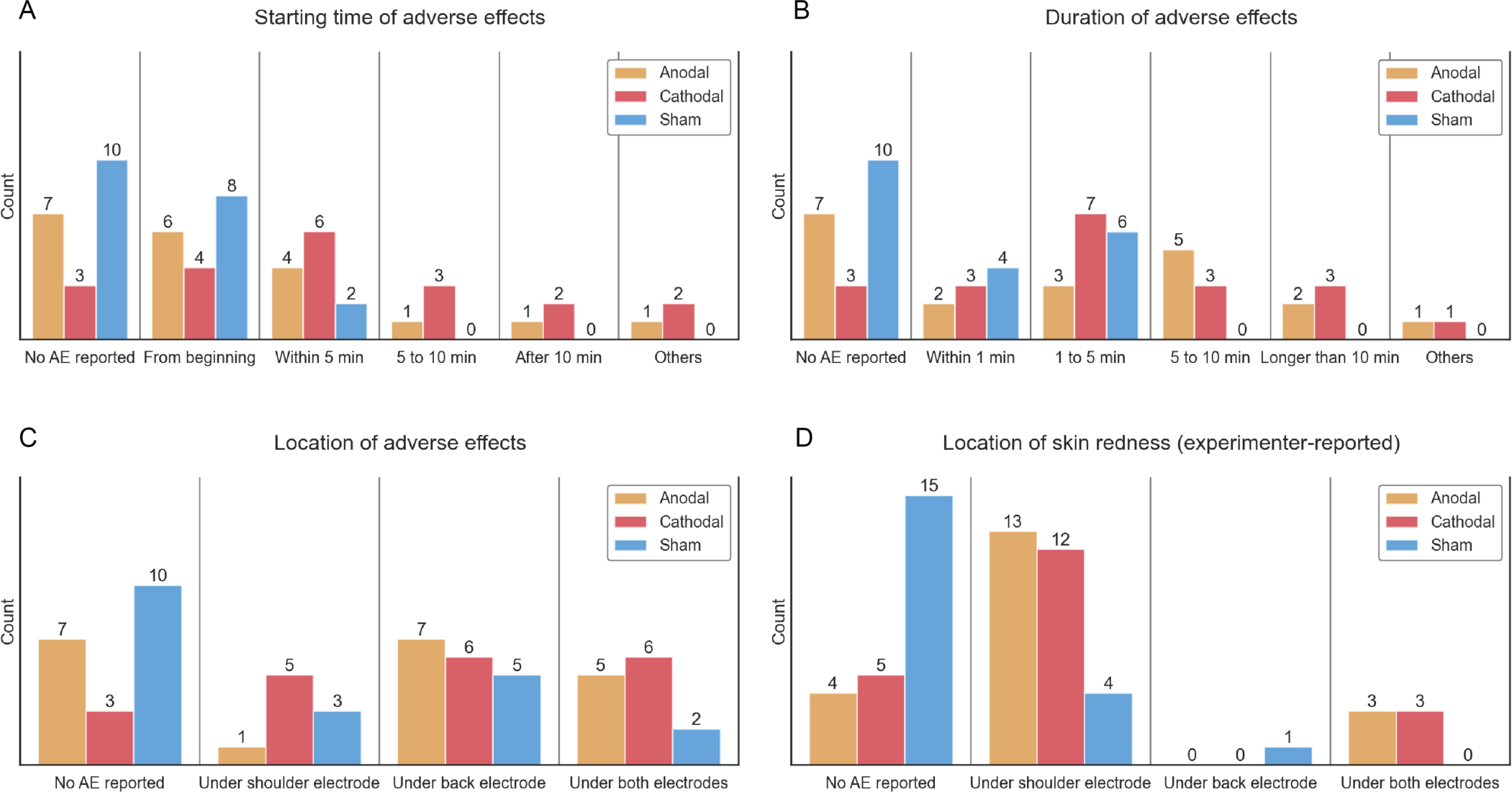
Temporal and spatial adverse effects patterns. Depicted reports of onset time (A), duration (B), location (C) of participant-reported AEs, and location (D) of experimenter-reported item across different the stimulation conditions. Please note that the category ‘Others’ was introduced as some participants reported differences in onset times and durations of AEs for electrodes, thus preventing an assignment to a unique category. Bars represent absolute numbers of reports among the sample of 20 participants.

### 3.3 Assessment of unspecific effects (UEs)

Participant-specific and group-level scores of the tsDCS-concurrent physiological measures are depicted in Figure 4. Out of the ten statistical comparisons, none showed significant differences and all BF were below one, with four instances providing moderate evidence for an absence of condition-differences (BF < 1/3; Table 4). To assess whether tsDCS-induced unspecific effects might have developed differentially over time, we tested for a time-by-condition interaction, but in eight out of ten statistical comparisons we did not observe significant interactions and in seven of those, BF provided moderate to strong evidence against an interaction effect (Table 5). Only for breathing rate did we observe a significant interaction, but the BF were equivocal and further investigation showed that this interaction was largely driven by a change of breathing rate in the sham condition (Figure 4F).

**Figure 4.**
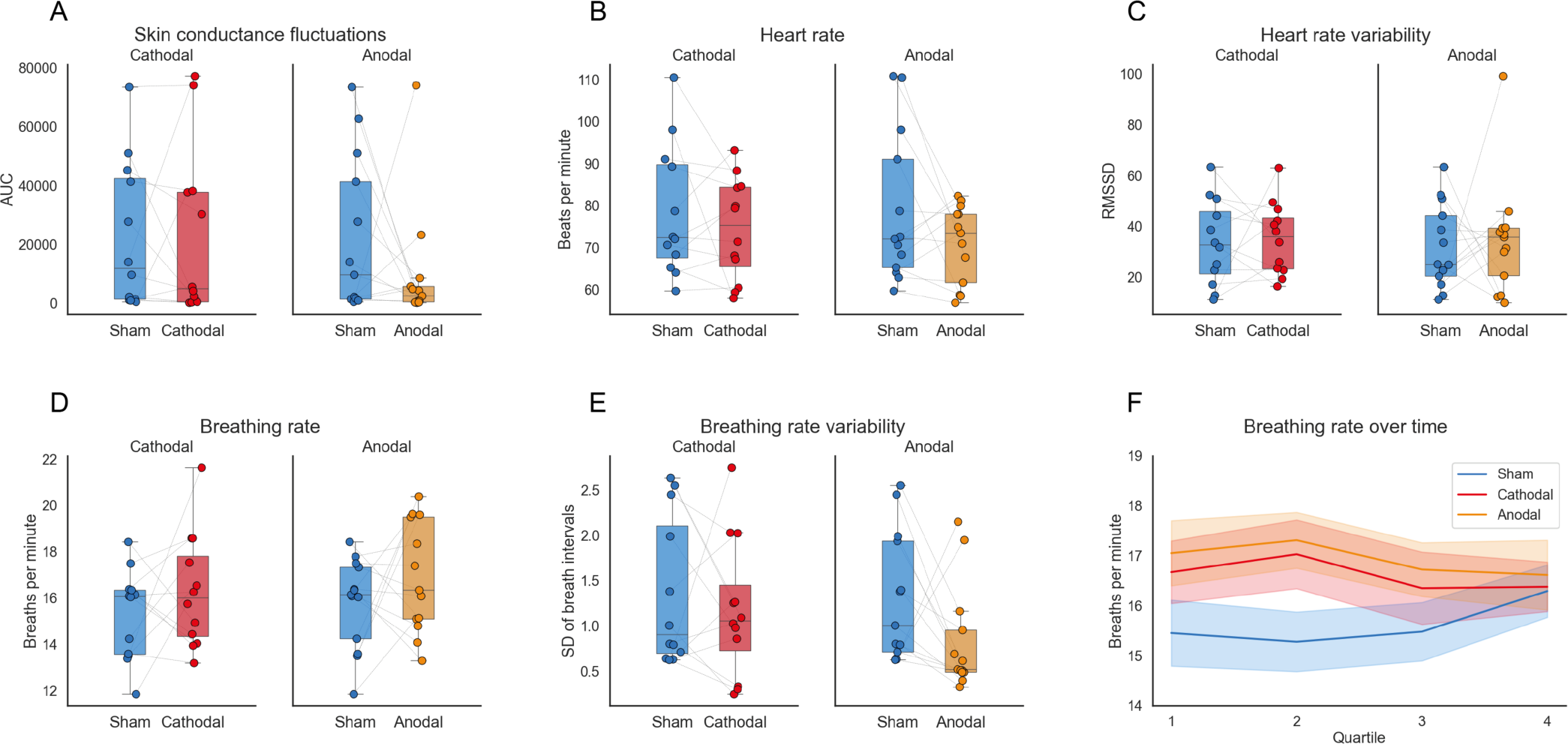
Unspecific effects assessment via autonomic responses in different stimulation conditions. Comparison of spontaneous skin conductance fluctuations (A), heart rate (B), heart rate variability (C), breathing rate (D), and breathing rate variability (E) between cathodal and sham as well as anodal and sham conditions, respectively. Note that the sham group does not always consist of the same data points as participants were excluded from specific sessions due to excessive noise (see description in Methods section). (F) Depiction of group-level means and standard error of the mean underlying the significant time-by-condition interaction in breathing rate.

**Table 4.**
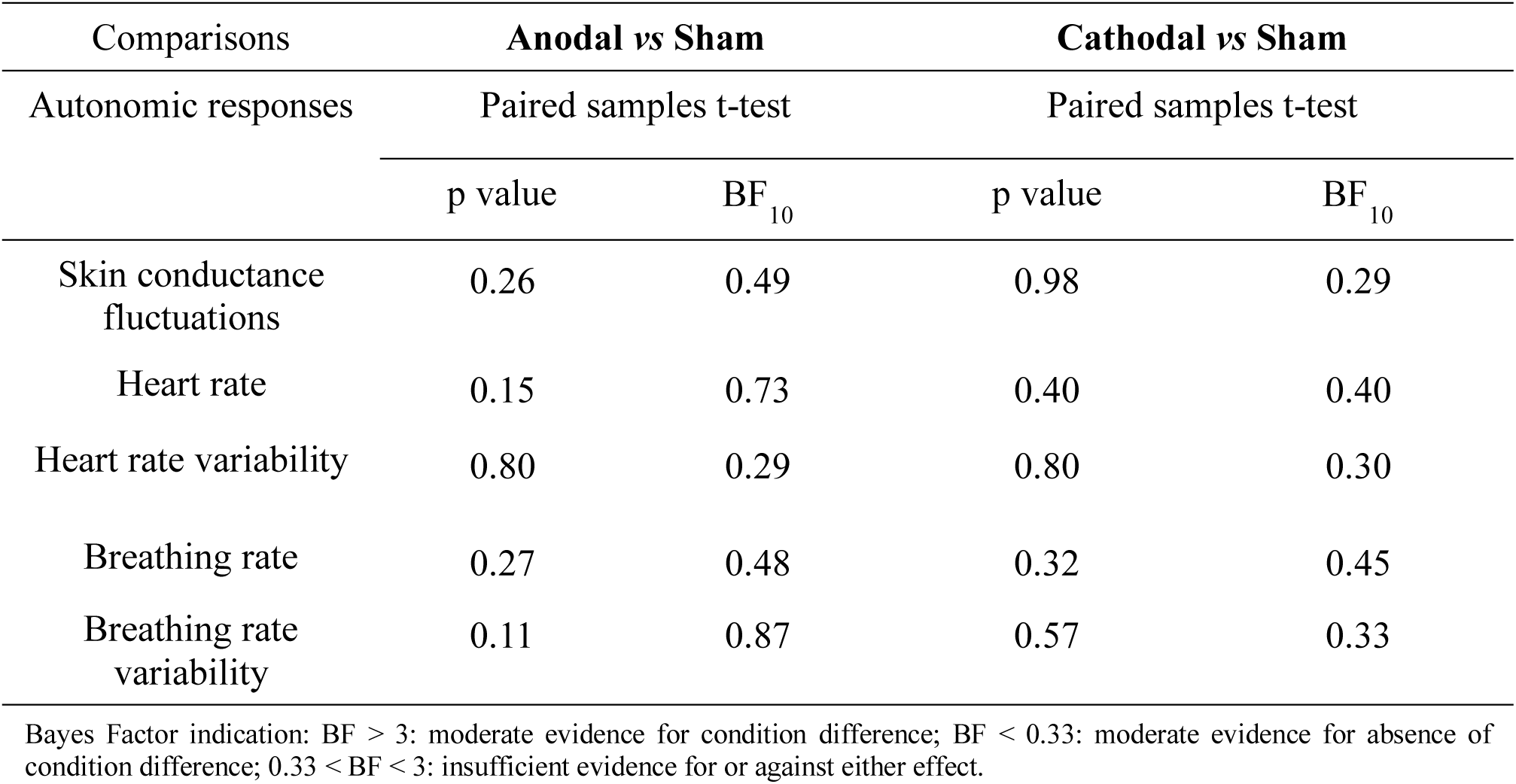
Statistical Comparison of UEs.

**Table 5.** Statistical Comparison of UEs – 5-minute intervals interaction effect.

| Comparisons | Anodal vs Sham |  |  | Cathodal vs Sham |  |  |
| --- | --- | --- | --- | --- | --- | --- |
| Autonomic responses | rm-ANOVA condition × time | | | rm-ANOVA condition × time | | |
|  | p value | BF<sub>incl</sub> | BF<sub>10</sub> | p value | BF<sub>incl</sub> | BF<sub>10</sub> |
| Skin conductance fluctuations | 0.21 | 0.16 | 0.08 | 0.26<sup>a</sup> | 0.08 | 0.04 |
| Heart rate | 0.74<sup>a</sup> | 0.02 | 0.01 | 0.27<sup>a</sup> | 0.10 | 0.05 |
| Heart rate variability | 0.85 | 0.05 | 0.03 | 0.63<sup>a</sup> | 0.05 | 0.02 |
| Breathing rate | 0.02 | 0.77 | 0.37 | 0.02 | 2.76 | 1.50 |
| Breathing rate variability | 0.23 | 0.11 | 0.06 | 0.09<sup>a</sup> | 0.95 | 0.42 |

BF<sub>incl</sub> indicates the change from prior to posterior inclusion odds (referring to the sum of the prior or posterior probabilities of all models that include the effect).
BF<sub>10</sub> indicates the comparison of our interaction model (including the two main effects) with a null model (containing only subject and random slopes).
Bayes Factor indication: BF > 3: moderate evidence for the tested effect; BF < 0.33: moderate evidence against the tested effect; 0.33 < BF < 3: insufficient evidence for or against either effect.
<sup>a</sup>Greenhouse-Geisser-correction used.

## 4. Discussion

Here, we investigated AEs and UEs associated with tsDCS, by first performing a review of the tsDCS literature in this regard and then empirically assessing AEs and UEs in a preregistered study via a structured questionnaire and tsDCS-concurrent physiological recordings, respectively.

### 4.1 Adverse effects (AEs) of tsDCS

To comprehensively assess tsDCS AEs in a structured way, we developed a questionnaire based on an existing tDCS template [16] and employed this in a randomized, within-participant, double-blind design involving 20 participants (who underwent anodal, cathodal, and sham tsDCS). This allowed us to provide detailed descriptions of overall AE reports as well as condition-differences using Frequentist and Bayesian statistics, including spatio-temporal AE aspects and blinding success. To our knowledge, this combination of factors goes far beyond what has previously been carried out in the tsDCS literature: out of 76 human tsDCS studies, only nine [24–32] employed structured questionnaires, with only three of these statistically comparing effects under active and sham conditions [25, 28, 29] and none investigating spatio-temporal aspects. Our study thus provides a starting point for a systematic and comprehensive assessment of tsDCS AEs, and we believe that the tsDCS community might benefit from a standardized and psychometrically evaluated questionnaire.

Our findings revealed predominantly mild AEs, mostly consisting of skin-related sensations at the electrode sites, such as burning, tingling, and itching. While this is in line with prior reports in the tDCS [17] as well as tsDCS literature [10, 12, 26, 33, 34], we went beyond these previous reports by conducting both frequentist and Bayesian comparisons between active and sham conditions for each AE: in none of the comparisons did we observe a significant difference on any item and complementary Bayesian analyses provided moderate evidence for an absence of condition differences in half of these comparisons. There were no reports of painful sensations or acute mood changes, which is in line with reporting in the tsDCS literature, where – across almost 80 studies – head pain [27] and musculoskeletal pain [35] were each only reported once; we furthermore observed only very few reports of sleepiness and trouble concentrating (rated as unlikely to be related to tsDCS). Taken together, this suggests that – from the perspective of participant-reports – tsDCS is a well-tolerated and safe technique, consistent with previous reports on the safety of tDCS [36].

We also asked participants about the onset, duration, and location of experienced AEs and observed that there were temporally more-extended AE reports as well as more reports of sensations under both electrodes in the active conditions. While previous tsDCS studies mostly focused on the presence or absence of AEs [29, 31], we believe that a spatio-temporal characterization of AEs as carried out here is important for allowing to design an appropriate tsDCS control condition that ensures adequate blinding.

### 4.2 Participant and experimenter blinding

Apart from AEs, we also investigated participants’ assumptions regarding the type of stimulation they received. While 50% of participants correctly reported that two sessions were active – suggesting their attentiveness to instructions [37], considering that this information was provided at experiment start and also upon questionnaire administration – only 5% were correct in assigning all three conditions, suggesting good blinding performance. While such a lack of correct condition assignment is in line with previous tsDCS studies [7, 11, 13, 14, 23, 24, 38–40], we went beyond this simple dichotomy and also explored whether participants’ accuracy in reporting the stimulation condition was associated with differences in reported AEs: reassuringly, also here we did not observe significant differences, suggesting that adequate blinding on the participant-side occurs even with a tsDCS intensity of 2.5mA as carried out here.

It is important to consider however that the experimenter-assessed item of skin redness clearly differentiated between active and sham conditions, potentially leading to experimenter-unblinding in the worst case [41]. Contrary to our observations (where skin redness was the most prominent AE), skin redness was only reported four times in the tsDCS literature [26, 42–44], without reports of significant differences between active and sham conditions as observed here, thus deserving further study. Another aspect to consider is how skin redness evolves over time, as participants could potentially unblind themselves regarding active vs sham stimulation by looking at their back / shoulder after the experiment.

Overall, we believe that it is prudent to formally assess blinding success regularly in tsDCS studies as well as investigate other approaches to sham stimulation, such as different electrode placement or expectation manipulation via de-facto masking [41, 45].

### 4.3 Unspecific Effects (UEs) of tsDCS

We also assessed whether active compared to sham tsDCS induces UEs in bodily state and observed consistently non-significant results as well as Bayes Factors mostly indicating an absence of condition-differences. This pattern of results suggests that active thoracolumbar tsDCS does not modulate vital functions such as heart rate or breathing rate, which is reassuring from a safety perspective. Despite our findings, we believe that further research is necessary to replicate and extend these results, considering that our systematic review indicated that this field is virtually untouched: one study investigated longitudinal post-tsDCS changes in skin conductance (though in patients where an autonomic dysfunction is part of the pathology) [32] and another study assessed changes in spontaneous breathing as well as skin conductance and heart rate (though the latter two not in a polarity-dependent or sham-controlled manner) [23].

The absence of effects on autonomic function observed here is also noteworthy when considering the spatial proximity of our stimulation site (12^th^ thoracic vertebra) to some of the autonomic outflow pathways. The sympathetic nervous system originates from the T1 to L3 levels of the spinal cord [46, 47], with a focus on T1 to T5 for upper limb and cardiac innervation. Modelling studies exploring the E-field of thoracolumbar tsDCS [48–50] suggest that such thoracic segments could be affected by our type of tsDCS. Conversely, the phrenic motoneurons innervating respiratory muscles are located in the spinal segments C3–C5 [51], which should not be affected by our type of tsDCS. Taken together, we believe that the tsDCS community should routinely record autonomic signals during experiments, as these are easy to obtain and would offer important insights into tsDCS’s specificity and safety.

### 4.4 Limitations and future directions

Several limitations of our study are worth mentioning. First, our AE and UE assessment occurred in young healthy volunteers and thus has limited generalizability to other populations. Second, a more comprehensive exploration of bodily states (including metrics such as blood pressure and cortisol levels) would offer a more holistic understanding of the off-target impact of tsDCS. Third, our focus on the acute effects of tsDCS does not allow any inferences on the cumulative effects of repeated tsDCS sessions, as would be relevant clinically. Fourth, our results only pertain to thoracolumbar tsDCS and it is thus essential to carry out similar studies for cervical tsDCS (which might have different UEs). Finally, our results suggest that maintaining experimenter and participant blinding requires considerable attention in future studies and possibly also more sensitive assessments of blinding success than the here-employed “end-of-study guess” [52].

## 5. Conclusion

Our investigation into the AEs and UEs of tsDCS demonstrates that tsDCS is a safe and well-tolerated technique, whose AE profile is primarily characterized by mild skin-related effects. Our UE findings furthermore indicate that tsDCS does not cause alterations in core autonomic measures and could thus be expected to exert rather specific neural effects. Taken together, our study provides substantial contributions to the understanding of tsDCS safety and specificity as well as participant blinding and should be followed up by similar approaches in clinical populations and longitudinal studies to unlock the full potential of tsDCS for understanding and modulating spinal cord function in health and disease.

## CRediT authorship contribution statement

**Hongyan Zhao**: Conceptualization, Data curation, Formal analysis, Investigation, Methodology, Project administration, Software, Visualization, Writing – original draft, Writing – review & editing. **Ulrike Horn**: Data curation, Formal analysis, Methodology, Software, Visualization, Writing – original draft, Writing – review & editing. **Melanie Freund**: Project administration, Writing – review & editing. **Anna Bujanow**: Resources, Writing – review & editing. **Christopher Gundlach**: Methodology, Writing – review & editing. **Gesa Hartwigsen**: Methodology, Writing – review & editing. **Falk Eippert**: Conceptualization, Funding acquisition, Methodology, Project administration, Resources, Supervision, Visualization, Writing – original draft, Writing – review & editing.

## Declaration of competing interest

The authors declare that they have no competing financial interests or personal relationships that could have appeared to influence the work reported in this manuscript.

## Acknowledgements

The authors would like to thank Magdalena Gruner, Josephine Graefe, Alicia Diem, and Mareike Pauly for their contribution to data collection and ensuring the rigorous completion of the experiment under a strict double-blind design as well as all volunteers for taking part in this study. This work was funded by the Max Planck Society and the European Research Council (under the European Union’s Horizon 2020 research and innovation programme; grant agreement No 758974).

## Data and code availability

The underlying data are openly available (https://osf.io/f7spw/; note to preprint readers: the data are currently only available to reviewers) as is all analysis code (https://github.com/eippertlab/tsdcs-sideeffects).

## Supplementary Material

**Supplementary Figure 1.**
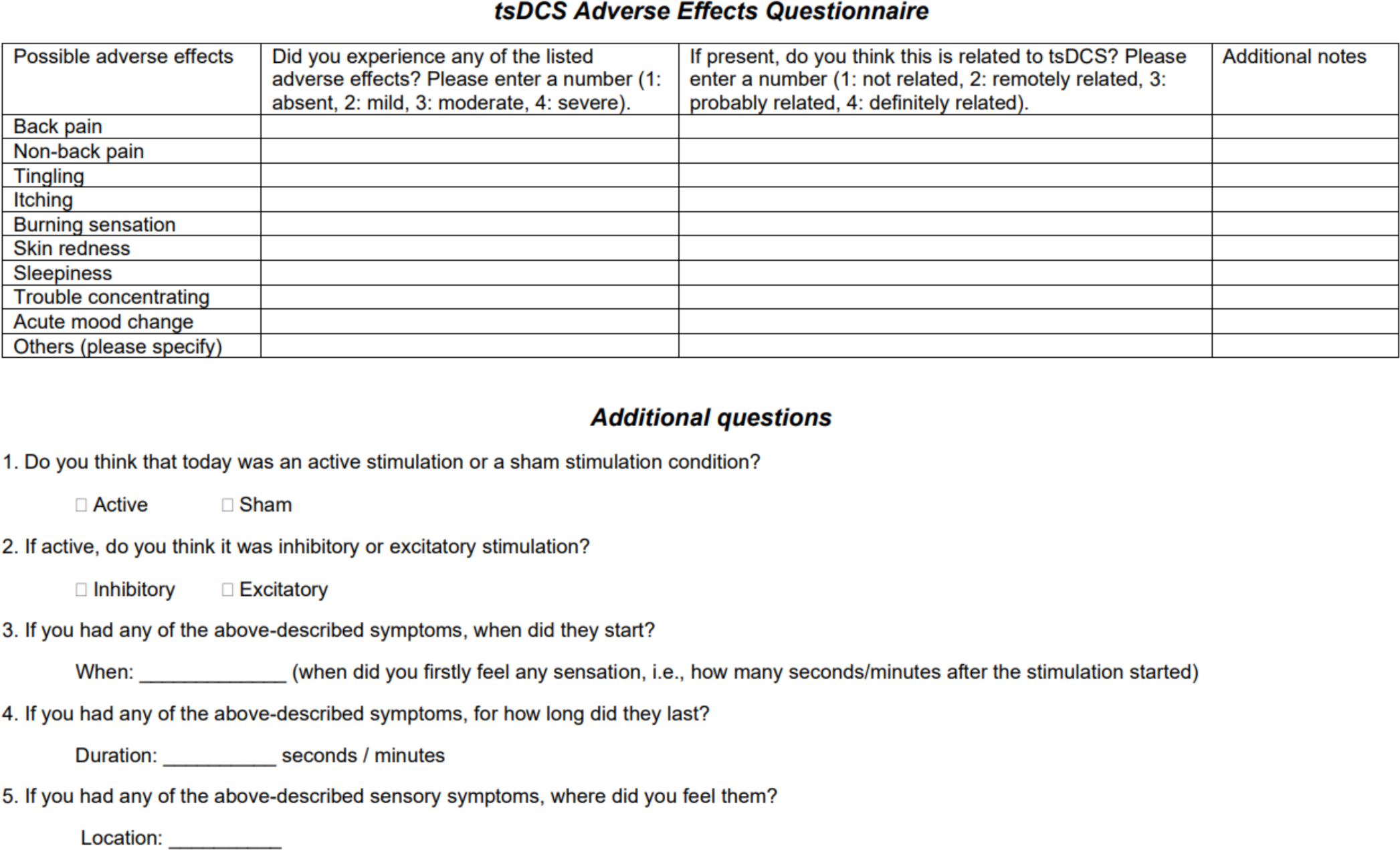
tsDCS Adverse Effects Questionnaire. The questionnaire, developed based on a proposed template for tDCS (Brunoni et al. 2011), captures potential adverse effect symptoms, their relation to tsDCS, participant guesses regarding the tsDCS condition, and details on adverse effects’ onset, duration, and location.

**Supplementary Figure 2.**
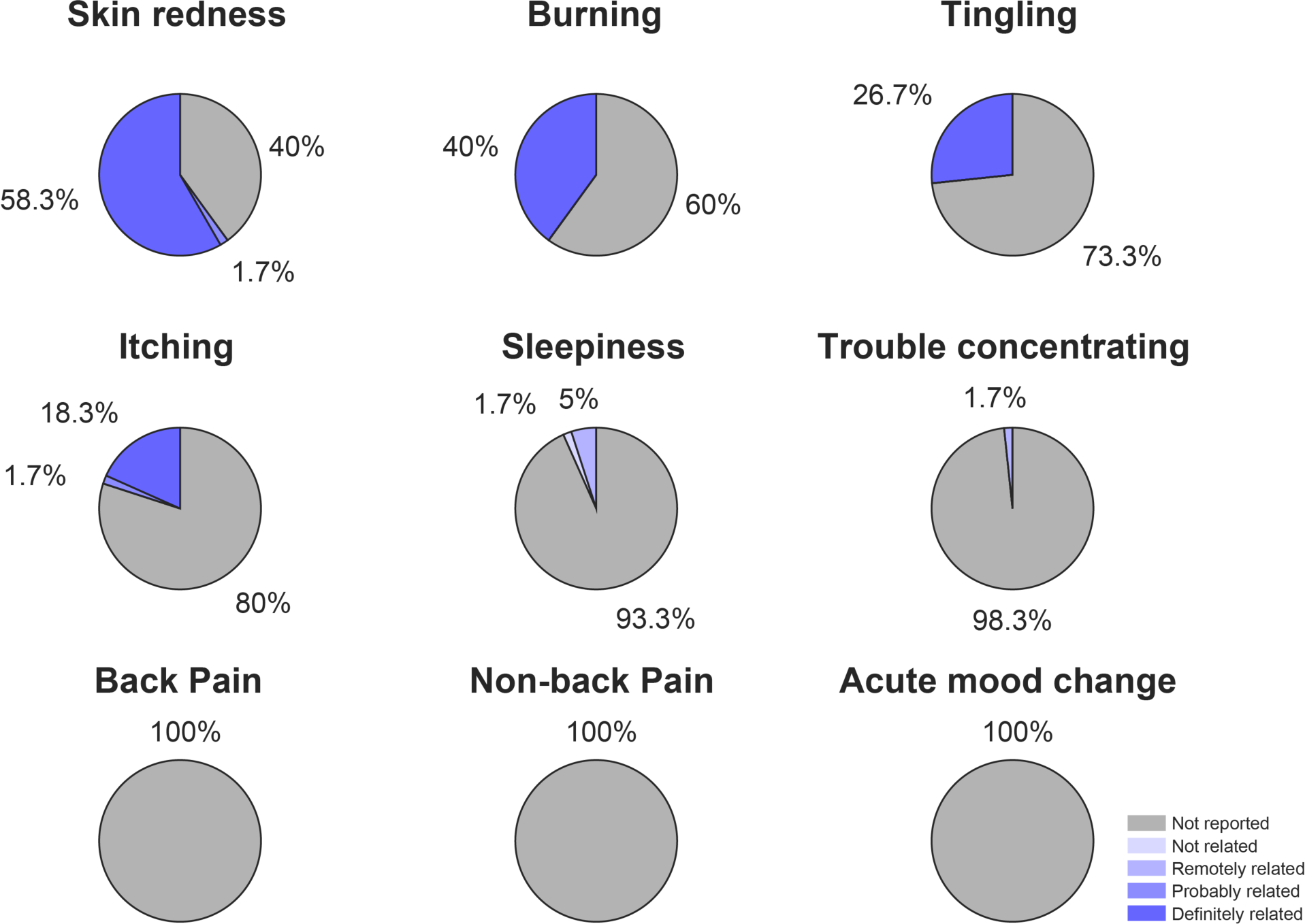
Relation reports of adverse effects with tsDCS. The relation of reported AEs is based on all 60 sessions, with colors representing the relation degree (see legend).

**Supplementary Table 1.**
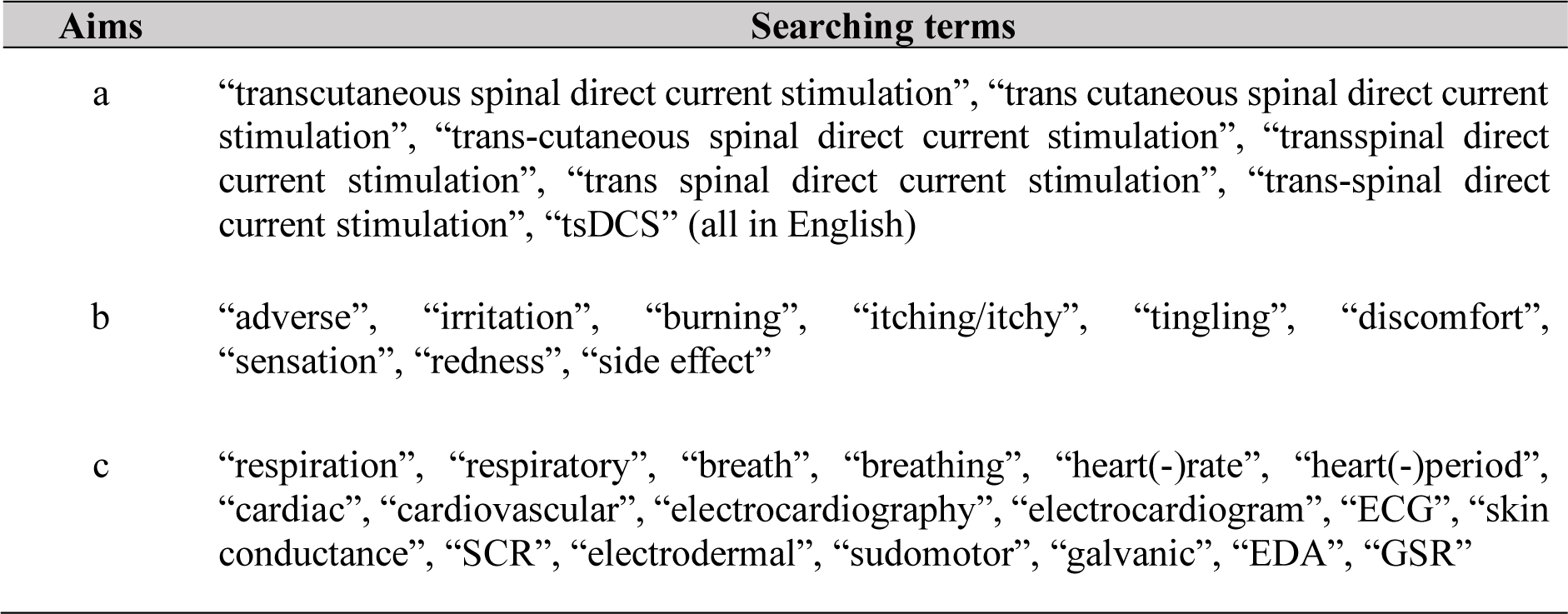
Searching terms for a) identifying studies, b) identifying the reporting of AEs and c) UEs.

**Supplementary Table 2.** Details on participant exclusion for UE analyses.

| Participant number | Session number | Reason |
| --- | --- | --- |
| 01 | 2 | talking |
| 08 | 2 | coughing |
| 09 | all sessions | excessive movement and talking |
| 10 | 3 | movement |
| 13 | all sessions | talking |
| 14 | all sessions | talking |
| 16 | 3 | talking |
| 17 | 3 | talking |
| 19 | 1 | movement |

**Supplementary Table 3.**
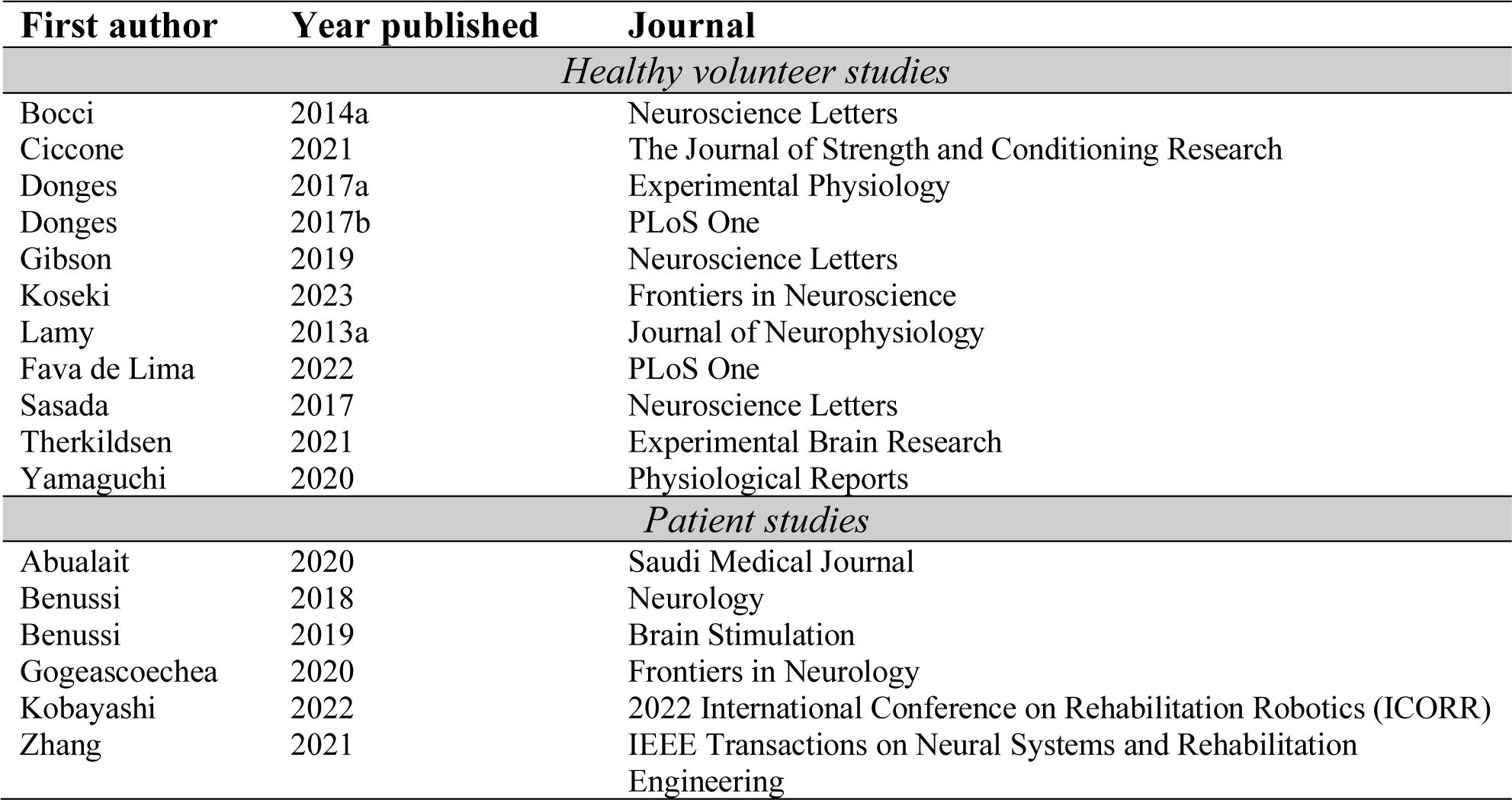
List of all studies in which our keyword search for tsDCS AEs did not return any hits.

**Supplementary Table 4.**
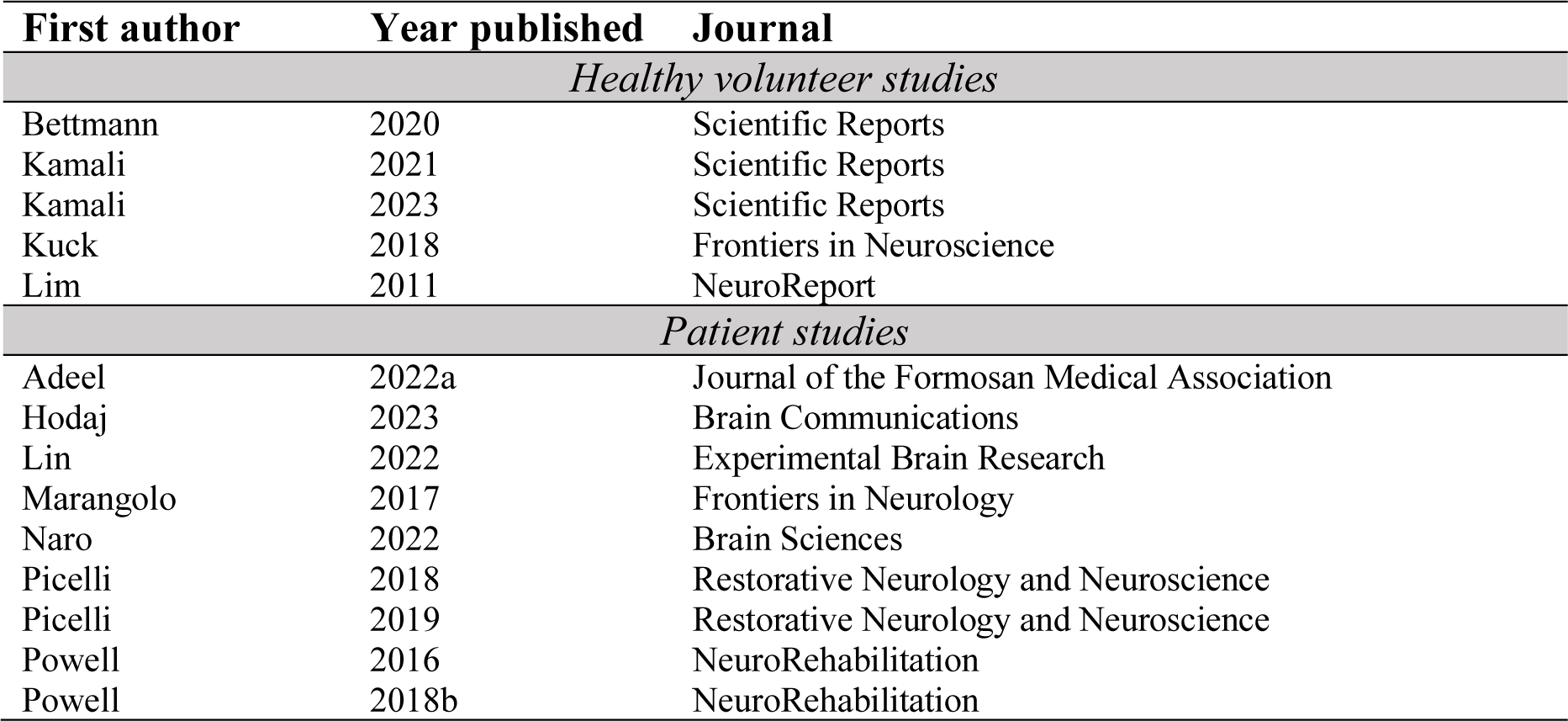
List of all studies in which our keyword search for tsDCS AEs did return hits, but where no AEs were reported.

## Notes

### Competing Interest Statement

The authors have declared no competing interest.

